# Show me your neighbour and I tell what you are: fisheye transformation for deep learning-based single-cell phenotyping

**DOI:** 10.1101/2022.08.23.505056

**Authors:** Timea Toth, Farkas Sukosd, Flora Kaptas, David Bauer, Peter Horvath

## Abstract

Recently we have concluded that image-based features derived from the microenvironment have an enormous impact on successfully determining the class of an object^1^. Here we demonstrate that deep learning-based phenotypic analysis of cells with a properly chosen microenvironment-size provides results comparable to our earlier neighbourhood-based methods that utilise hand-crafted image features. We hypothesised that treating cells with equal weight, regardless of their position within the cellular microenvironment, is suboptimal, and direct neighbours have a larger impact on the phenotype of the cell-of-interest than cells in its larger proximity. Hence we present a novel approach that (1) considers the fully featured view of the cell-of-interest, (2) includes the neighbourhood and (3) gives lesser weight to cells that are far from the cell. To achieve this, we present a transformation similar to those characteristic for fisheye cameras. Such a transformation satisfies all the above defined criteria, with a fast rate of transform for any images. Using the proposed transformation with proper settings we could significantly increase the accuracy of single-cell phenotyping, both in case of cell culture and tissue-based microscopy images. The range of potential applications of the proposed method goes beyond microscopy, as we present improved results on the *iWildCam 2020 dataset* containing images of wild animals.

## Introduction

All living entities adjust to their environment, manifested as visually observable morphological differences both at the macro- and microscales. Therefore, incorporating microenvironmental information into object classification may have an enormous impact on the accuracy of evaluation^1^. This phenomenon may be of particular interest when the phenotypes of cells are determined.

The modern technological advancements in microscopy, sequential hybridization^2,3^, and mass spectrometry^4^ have paved the way to evaluate cellular structures at high spatial and temporal resolution. These measurements generate large datasets, hence automated computational methods are required to obtain objective information from these images^5,6^. Utilising the inherent potentials of automated image analysis offers several advantages: it eliminates operator bias, provides quantitative data, and identifies visual characteristics that would otherwise go undetected.^7–9^

Various computer vision and classical machine learning techniques have been used^10^ to support researchers with tasks like image exploration (e.g. to find changes in cell structure in an imaging-based drug screen^11^), image classification (e.g. to determine the distribution of different proteins within cells^12^), image segmentation (e.g. identifying single cells in images^13^), or object tracking^14^. Despite the acknowledged capabilities of these techniques, deep learning-based analyses often perform more efficiently in recognizing biological patterns based on the pixels of images^15,16^.

Deep learning has yielded fascinating results in solving biology-related issues^17^. The phenotype of a cell is determined by various cellular processes and factors (including the stochasticity of gene expression, as well as a variety of proteomes and metabolomes^4^) that result in a particular morphological arrangement. Deep learning has enabled the exploration of factors like replicative age, organelle inheritance and response to stress^18^. It has been demonstrated to perform comparable to human pathologists upon classifying whole-slide images into two categories of cancerous and normal lung tissues (it was even able to predict the ten most commonly mutated genes)^19^. Another frequent challenge in cell biology lies in identifying various proteins and determining their locations within the cells. Numerous models have been developed^20,21^ to automatically identify subcellular localization patterns, based on the Human Protein Atlas^22^ which contains acquisitions of 12,003 human proteins at the single-cell level.

Single cell heterogeneity within cell populations is also influenced by the cell’s microenvironment^23,24^. Several studies have demonstrated that the peculiarities of cellular neighbourhood can be exceptionally relevant when investigating the collective organisation of cells in a variety of settings. Snijder *et al*. have reported that in a cell culture context one may predict the burden of viral infection at the single cell level, solely based on each cell’s microenvironment^24^. In a study of competitive interactions between wild-type Madin-Darby canine kidney (MDCK) cells and cells depleted of the polarity protein scribble, Bove *et al*. found that the probability of cell division is significantly higher for MDCK cells when their neighbourhood is mostly populated by scribble cells^25^. Several other examples of the importance of cellular neighbourhood are also published in literature. For instance, neighbouring epidermal stem cells affect each other (differentiation of a single stem cell is followed by division of a direct neighbour)^26^; ligand-producing hair cell precursors in the inner ear are smaller than their neighbours^27^.

In a previous work our group has also concluded that incorporating the features of the microenvironment of cells improves phenotype classification in high content screens^1^. In that study, we extracted commonly used cell-based features for every segmented cell. The centre of mass was measured for each segmented area and was used as a reference point for distance calculation. We used two different approaches to define cellular neighbourhoods: the K-nearest neighbours (KNN) and the N-distance methods (Fig. 1a). Neighbourhood features were derived from the mean, median, standard deviation, minimum, and maximum statistics of the previously calculated cellular features. Then, we used these neighbourhood features to classify cells and we got the best result using Multi-layer Perceptron classifier. Based on these findings, we hypothesised that it is worth using environmental data for deep learning phenotypic profiling.

**Figure 1.**
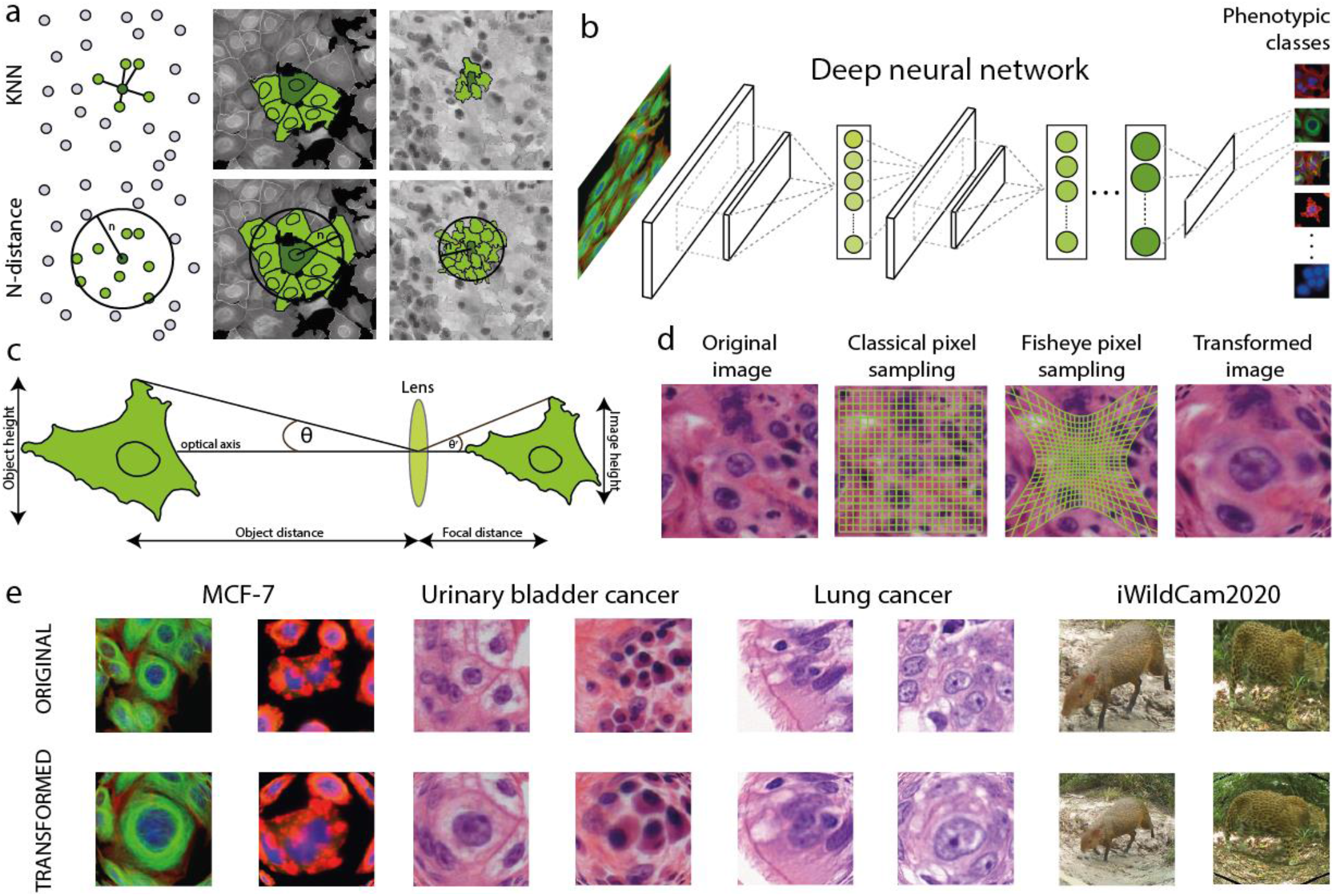
(a) Neighbourhood with classical machine learning^1^. The K-nearest neighbours (KNN) and N-distance methods, illustrated in a schematic figure and in real cell culture and tissue section scenarios (K=5, cell culture: n=19.51 μm, tissue sections: n=13.5 μm). (b) Schematic figure of phenotypic classification with deep learning. (c) Illustration of the optical parameters for the fisheye transformation. (d) The difference between classical and fisheye pixel sampling: in the classic case we select pixels evenly, while in case of fisheye, sampling is dense near to the object-of-interest, and less dense as the distance from the object increases. (e) Examples of the fisheye transformation

Recently, fisheye cameras have received significant interest from both technical professionals and the public in general. Fisheye lenses are ultra-wide angle lenses capable of taking wide panoramic or hemispheric images, however, they incorporate a significant optical distortion into the process. Fisheye lenses utilise specific mapping (stereographic, equidistant, equisolid angle, orthogonal) which lands the images a characteristic convex non-rectilinear appearance^28^. These specific lenses have a wide range of applications due to their ability to provide rich visual information, including the generation of augmented or virtual reality^29^, improving the performance of intelligent robot vision systems^30^, and simplifying the complexity of surveillance systems^31^. Various correction models have been proposed to rectify the distortion of fisheye lenses^32–34^.

In this paper we introduce a novel way of representing images to deep learning-based image classification networks. The basic idea is the following: the original image includes the object of interest (which is located in the middle of the image), as well as its microenvironment of a predefined range. The images are then transformed by a fisheye-like spatial sampling method that collects more pixels from the close proximity of the object-of-interest, and the resolution decreases for larger proximity (Fig. 1d). Our results indicate that the proposed transformation highly outperforms classical machine learning methods and deep learning-based classifiers benchmarked on cell cultures, scans of cancerous tissues and real life images. We remark that our fisheye transform method provides improved classification accuracy, as demonstrated by higher accuracy scores on several test datasets, compared to relying on feeding the network with multi-scale images in parallel (i.e. an image pyramid). Also, the presented fisheye transformation has the advantage that it can be incorporated into the network as a layer^35^, although technically it is more resource intensive when large images are fed into the network.

## Materials and methods

### Datasets

#### MCF-7 High-Content-Screening Dataset

We used a publicly available breast cancer cell line (MCF-7) dataset that included images of cells treated with 113 different small molecules at eight different concentrations for 24 hours (available online at the Broad Bioimage Benchmark Collection https://www.broadinstitute.org/bbbc/BBBC021/). In brief, the MCF-7 cell line was treated with a variety of targeted anticancer agents and standard cytotoxic compounds, resulting in a wide range of visible and subtle phenotypic changes at the cellular level. After 24 hours of incubation, the cells were fixed and stained for DNA, F-actin and B-tubulin before being scanned by fluorescence microscopy. Images were collected from 55 microtiter plates of 96-well format. The dataset contains more than 39,000 images of almost 2 million cells^36^. Piccinini and colleagues^37^ published a single-cell phenotypic annotation method which we have adopted in our study. Nine phenotypic classes (i.e. abundant, rounded, elongated, multinucleated, bundled microtubule, peripheral cytoskeleton, punctate actin foci, decreased cell size and fragmented nucleus) and a debris class were identified (Fig. 2a), and approximately 1,500 cells were labelled.

**Figure 2.**
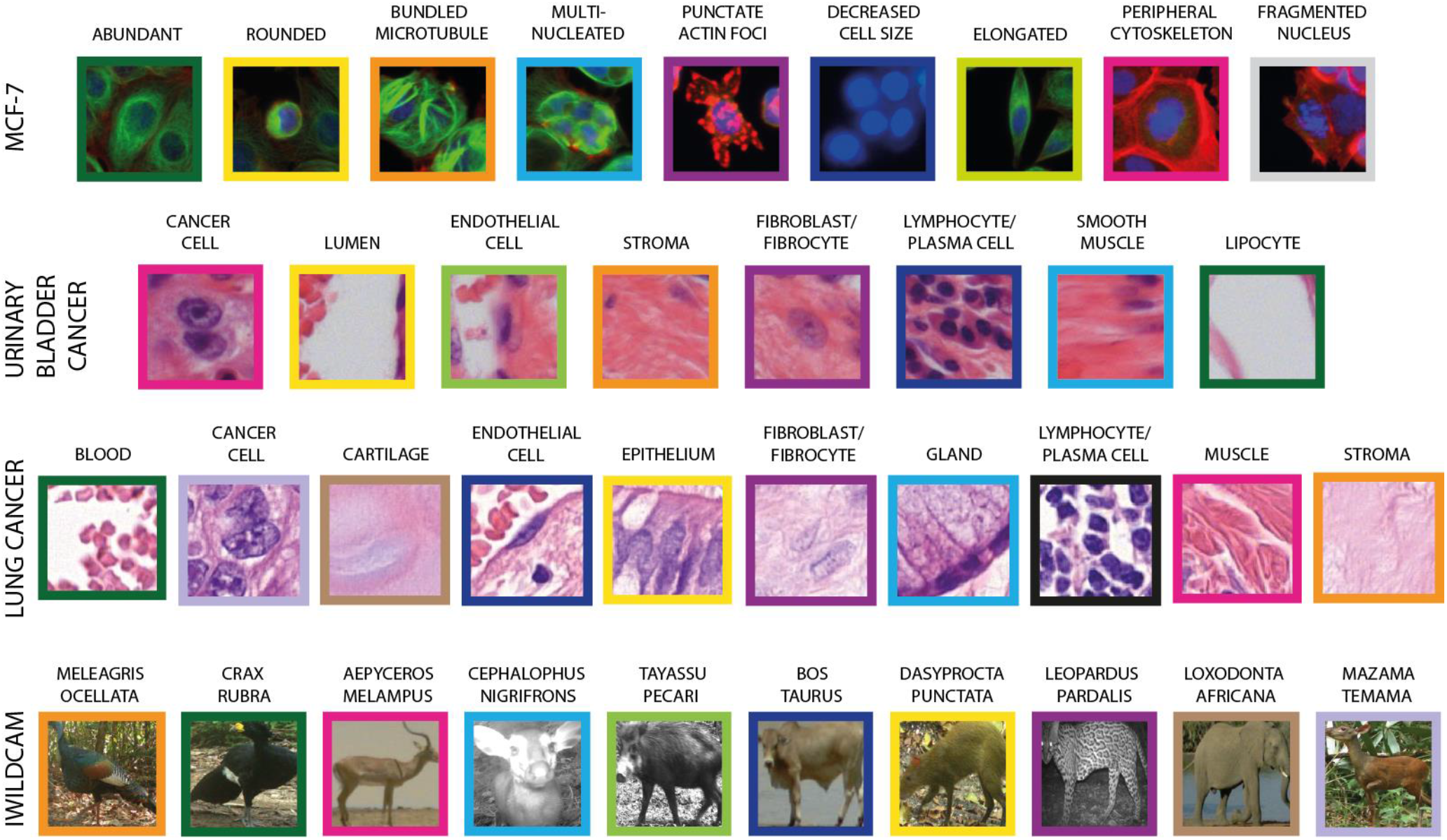
Distinguished classes. (a) Cells of nine different phenotype classes identified in the MCF-7 High-Content-Screening Dataset. (b) Eight phenotypic classes in the UBC tissue image dataset. (c) Ten phenotypic classes in the LC tissue image dataset. (d) The ten most common animal species in the iWildCam2020 dataset.

#### Urinary bladder cancer and lung cancer tissue sections

Images of urinary bladder cancer (UBC) and lung cancer (LC) tissues served as our second and third test datasets (Fig. 2b,c). The slides of urinary and lung cancer tissues were stained with hematoxylin-eosin (HE) in standard histopathological procedures. Formalin-fixed and paraffin-embedded tissue sections were cut into 4 µm thick slices, and were stained using a Tissue-Tek DRS 2000E-D2 Slide Stainer (Sakura Finetek Japan) according to the manufacturer’s instructions. Using the AxioVision SE64Rel.4.9.1.1 (Carl Zeiss Meditec AG, Germany) software, images were captured with an Axio Imager Z.1 (Carl Zeiss Meditec AG, Germany) microscope equipped with an EC Plan-NEOFLUOAR 20x/0.5NA lens.

For the analysis of the bladder cancer tissue dataset, containing 38 images, we used the annotation presented in our previous neighbourhood study^1^. We distinguished eight phenotypic classes (cancer cell, lumen cell, endothelial cell, stroma cell, fibroblast-fibrocyte, lymphocyte-plasma cell, smooth muscle cell and lipocyte), and labelled 1,200 cells. For the lung cancer dataset, we differentiated 10 phenotypic classes (blood cell, cancer cell, cartilage, endothelial cell, epithelium, fibroblast-fibrocyte, gland, lymphocyte-plasma cell, muscle cell and stroma cell), and labelled 5,000 cells.

#### iWildCam 2020 dataset

The iWildCam 2020 dataset is derived from the iWildCam 2020 competition organised by Kaggle, where the competition task focused on classifying animal species. Data were primarily provided by the Wildlife Conservation Society, iNaturalist, the U.S. Geological Survey, and Microsoft AI for Earth. The training dataset of the competition consists of 217,959 images taken at 441 locations. In total, 267 classes, containing a highly unbalanced number of entities, were defined. Fig. 2d displays the 10 classes that contain most of the elements (excluding the ‘empty’ class, where no animals are visible in the image).

### Segmentation

CellProfiler 2.2.0 was used to segment images from the high-content-screening dataset of drug-treated MCF-7 samples. The adaptive Otsu method was used to detect nuclei. Cells smaller than 5 µm and nuclei contacting the borders of the images were eliminated. Adaptive thresholding was used to extract the cytoplasm of cells, using watershed separation based on the nuclei as seed points.

For the segmentation of the tissue datasets and for the annotation of the MCF-7, UBC and LC datasets we used an image analysis and machine learning software named BIAS^38^, which was developed by Single-Cell Technologies Ltd. (Szeged, Hungary).

The SLIC superpixel segmentation technique^39^ was employed to segment urinary bladder cancer and lung cancer section images. We set 35 pixel as superpixel size, and forced connectivity between superpixels when a superpixel included less than 25 pixels. We have previously demonstrated that this superpixel size works best to represent the cellular structure of tissues^1^.

During the annotation process of these three (MCF-7, urinary bladder and lung cancer) datasets, we saved the x-y coordinates of the centre of the nuclei/superpixels, and used these coordinates as inputs for the fisheye transformation.

For the iWildCam dataset, the organisers of the competition provided a general animal detection model (https://github.com/microsoft/CameraTraps/blob/master/megadetector.md) called MegaDetector, along with an annotation file that contained one label per image. When more than one animal were visible in the image, we selected only the detection with the highest accuracy, and gave the label to it. MegaDetector works with bounding boxes. In the present study, we used the x-y coordinates of the centre of the bounding boxes as inputs for the fisheye transformation.

### Fisheye transformation

Several types of ultra-wide angle lenses are available, and all are associated with a significant visual distortion. In our study we evaluated an algorithm that artificially reproduces the same kind of distortion which is inherent in images taken with ultra-wide angle lenses. We analysed the relevance of neighbourhood features to determine the most optimal distance to be considered for the highest accuracy of classification.

The first parameter we optimised was the range around our object-of-interest (in an optical manner, this is object height, later referred as ‘window size’, see Fig. 1c). For the MCF-7, UBC and LC datasets, as we have mentioned before, we saved the x-y coordinates of the centre of the nuclei/superpixels and then we used the selected pixel range for the fisheye transformation based on these points. In case of the iWildCam dataset, we cropped out areas of interest from the original images for our fisheye transformation in four different sizes. We took the size of the original bounding box and multiplied it with 1.0, 1.5, 2.0 and 2.5. When we multiplied the original size with 1.0, we cropped the images with the same size for the baseline images and for the fisheye-transformation. In all of the other cases, we took a bigger size of the environment into account.

The second modifiable parameter was focal distance, which essentially defines the strength of the distortion. Contrary to our original neighbourhood idea^1^, information acquisition was not cell-based, but pixel-based (Fig. 1a,d). The third parameter was the mapping function, which originally (in cameras) is responsible for transforming a part of a spherical object to a 2D plane. In the present study we chose the ‘equidistant’ function -as it is one of the most popular mapping functions used in cameras - to test our hypothesis on the significance of neighbourhood regarding classification accuracy. In all cases we set the object distance (the distance between the original image and the lenses) in a way to eliminate scaling due to fisheye distortion. For more information about the transformation and examples of the transformed images, see Supplementary Section 1 and Supplementary Figure S1-S4.

### Deep learning-based object classification

For image classification we used Matlab R2019b and its Deep Learning Toolbox (version 13.0). This toolbox provides a framework to design or implement networks, pre-trained models, and apps. Two networks pre-trained on ImageNet, ResNet50 and InceptionV3 were utilised for transfer learning. Our decision for the models (i.e. using ResNet50 and InceptionV3), was based on the accuracy, speed and size of the networks. Particular attention was paid to avoid that an annotated cell is included in the training dataset and appears in a validation image as a neighbour afterwards. Thus, to prevent a potentially positive influence of evaluation, both the MCF-7 and UBC datasets were processed with pre-defined train and validation folders at the image level, instead of the cell level. For the LC and iWildCam datasets this was not an issue, as these were created to contain only one annotated object per image. The latter two datasets were randomised and split to a training and a test set, thus 80% was used for training and 20% was used for testing the methods. Three independent trials were run for each dataset.

Data augmentation has a seminal role in improving classification accuracy. We applied the following standard geometry transformations on our training datasets: reflection in the left-right and a top-bottom directions, rotation, horizontal and vertical scaling. It is important to mention that we have not used any transformation that spatially moves the cell-of-interest away from the centre of the image because that would have consequences to the fisheye transformation.

## Results

We evaluated the performance of a fisheye-like sampling on several image datasets, aiming to improve classification accuracy by deep learning-based image classification networks. We compared two convolutional neural network-based classifiers using ResNet50 and InceptionV3 backbones, as well as the relevance of the extent of neighbourhood and the focal distance of fisheye transformation. We benchmarked what combination of these parameters produce the best results.

The baselines we compared our method were also calculated with ResNet50 and InceptionV3 models. In case of the cell culture and tissue section datasets, the input images were generated as: based on the centre of the cells, we cropped out 192×192 pixel-sized images around them, then resized these images to 224×224 to meet the requirement of the models. For the iWildCam dataset, we cropped out the inside of the bounding boxes that were drawn around the animals, then resized these to 224×224 pixel images. For benchmarking, we have not used the fisheye transformation. For the statistical analysis of the classification accuracy results, two-sample t-tests were performed (Supplementary Section 4).

Based on previous studies in cellular biology, we expected that taking the cell’s microenvironment into consideration improves the performance of deep learning classification. To test this hypothesis, we have introduced a fisheye transformation, as this kind of distortion considers more pixels from the direct neighbourhood of the object-of-interest than from the region beyond that (Fig. 1d). We also hypothesised that there should be an optimal neighbourhood range and focal distance combination with respect to deep learning based classification accuracy.

### Increased classification accuracy on images of cell cultures

In case of the MCF-7 breast cancer cell database, an environmental range of 45 to 724 pixels (17.56-282.54 µm) was defined. For comparison, the average nuclei size in this dataset is 37 pixels (14.44 µm). Validation results indicate that applying the fisheye transformation improves the accuracy of both (Resnet50 and InceptionV3) classifiers (Fig. 3a). The best performance was achieved when we applied a window size of 543 pixels (211.91 µm) with a focal length of 130 arbitrary units (where 1 unit corresponds to the size of a pixel), using the ResNet50 model. In this case, accuracy reached 91.38%, which is 7% better than that achieved when using deep learning only (84.31%). The highest classification accuracy achieved with fisheye distortion also outperformed our previous results with classical machine learning approaches^1^, where the maximum accuracy was 90.80% with the support vector machine classifier. Although the deep learning baseline was higher for InceptionV3 (85.85%) than for ResNet50, the best result we could achieve with InceptionV3 using distorted images was 89.33% only.

**Figure 3.**
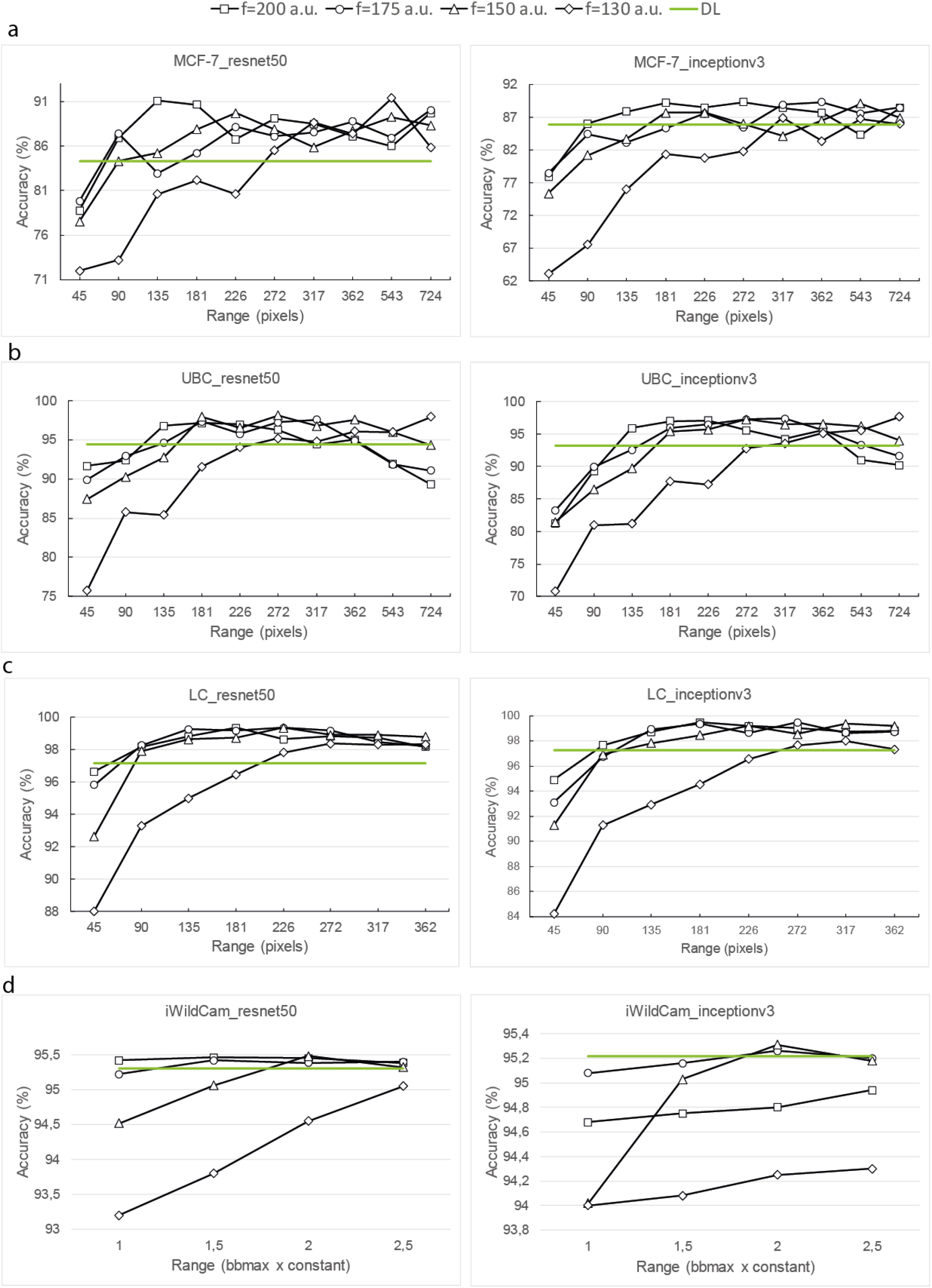
Comparison of the performance of deep learning networks (ResNet50, InceptionV3) upon considering different neighbourhood distances. (a) Classification accuracies for the MCF-7 cell culture dataset using ResNet50 (left) and InceptionV3 (right). (b) Classification accuracies for the urinary bladder cancer tissue image dataset. (c) Classification accuracies for the lung cancer tissue dataset. (d) Classification accuracies for the iWildCam2020 dataset. Green lines indicate the baseline yielded with deep learning upon using the original (undistorted) images, while black lines indicate the results achieved on fisheye distorted images with different f (focal distance) values (the values are measured in arbitrary units).

**Figure 4.**
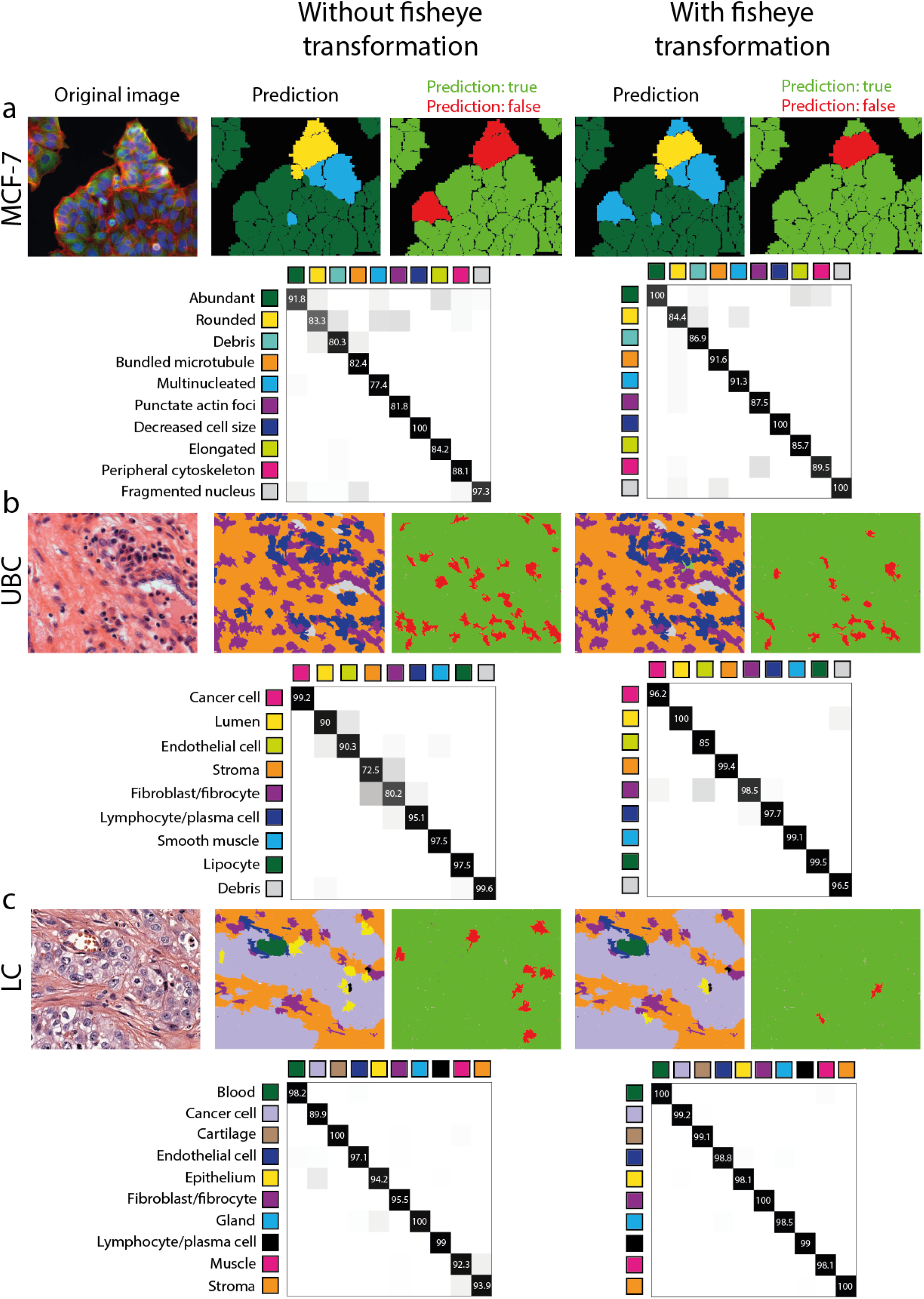
The effect of combining the fisheye transformation with deep learning. (a) Prediction examples and confusion matrices based on ResNet50 models in the cell culture dataset. Original image (left), prediction and confusion matrix using the model built on standard images (middle), prediction and confusion matrix using the model built on fisheye-transformed images, window size: 543 pixels, focal length: 130 a.u. (right). In the second and fourth images we show with different colours the predicted phenotypes and in the third and fifth images we show if the prediction was true (green) or false (red). (b) Prediction examples and confusion matrices of the best deep learning performance (ResNet50) in the urinary bladder cancer tissue dataset. Original image (left), prediction and confusion matrix using traditional deep learning (middle) and considering the cellular neighbourhood with fisheye transformation, window size: 272 pixels, focal length: 150 a.u. (right). (c) Prediction examples and confusion matrices based on InceptionV3 models in the lung cancer tissue dataset. Original image (left), prediction and confusion matrix using the model built on undistorted images (middle), prediction and confusion matrix using the combination of deep-learning and fisheye-distorted images, window size: 272 pixels, focal length: 170 a.u. (right).

In order to demonstrate that the increase in classification accuracy is due to the inclusion of the environment and the fisheye transformation, we performed additional experiments on images containing only the nucleus, or the nucleus and cytoplasm of the cell (see Supplementary Section 5). Both of these data largely underperformed the proposed fisheye model achieving only 80.33% in case of nuclei and 82.29% in case of whole cell images.

### Fisheye transformation has a major impact on phenotyping tissue sections

In case of the UBC tissue image dataset, we also collected neighbourhood information for a 45 to 724 window size (12.15-195.48 µm), where the average nuclei size is 32 pixels (8.64 µm), while for the lung cancer dataset we used a range of 45 to 362 pixels (17.55-141.18 µm) with average nuclei size 39 pixels (15.21 µm). For both datasets, we were able to achieve higher classification accuracy on the fisheye transformed images than with traditional deep learning, irrespective of whether we used ResNet50 or InceptionV3.

For the UBC dataset, we had previous results using neighbourhood features with classical machine learning^1^. Then the maximum of classification accuracy reached 93.37% with MLP (multilayer perceptron) calculations. In our current study the best performance reached 98.14%, appearing at a window size of 272 pixels with 150 arbitrary unit focal length using ResNet50 (Fig. 3b). Without using the distorted images, classification accuracy was 94.41% only. Using InceptionV3, both the deep learning baseline (93.24%) and the highest accuracy achieved on fisheye distorted images (97.65%) were less favourable than those yielded with ResNet50.

The lung cancer dataset is the only exception where InceptionV3 performed slightly better than ResNet50 (Fig. 3c). With ResNet50, the highest classification accuracy was 99.36%, while InceptionV3 yielded a maximum of 99.46% upon incorporating the neighbourhood feature with a 272 pixel range and using 170 arbitrary units as focal distance. This is more than 2% better than the results yielded with InceptionV3 on undistorted images (accuracy: 97.25%).

### Fisheye transformation outperforms image pyramids

In order to benchmark our proposed fisheye transformation against classical multiscale approaches, we tested the phenotypic classification accuracy on UBC and LC datasets using image pyramid as an input to the networks. For this purpose, we fed the ResNet50 networks with images of varying (1/1, 1/2, 1/4) scales in parallel. Accuracy reached 97.9% and 99.1% for the UBC and lung cancer datasets, respectively, indicating that applying the image pyramid approach yields better results than traditional deep learning, but performs less favourable compared to the fisheye distorted solution.

### Improved accuracy in case of the iWildCam2020 dataset

We investigated whether the inclusion of the environment of real scenes can improve classification results. The dataset included photos of animals taken from fixed camera positions. Because of the perspective, the animals could appear either very small or large in the images. To handle this sort of discrepancy, we considered the size of the bounding boxes around the animals as references, instead of using fixed pixel-distances. The baselines for classification accuracy with traditional deep learning were 95.3% and 95.22% upon using ResNet50 and InceptionV3, respectively. In case of fisheye-transformed images, classification accuracy reached 95.48% with ResNet50, when 2.5× the size of bounding boxes were considered as the neighbourhood feature and focal length was set to 150 units.

## Discussion

Here we present a method combining fisheye transformation with deep learning, an extension to our previous model, incorporating the information obtained from the cellular microenvironment in phenotypic classification. We demonstrated on MCF-7 cell culture and Urinary Bladder Cancer datasets, that our method outperforms the approach of using classical machine learning with single-cell and neighbourhood features. Our results were compared to the accuracies obtained with deep learning (ResNet50 and InceptionV3), where the input for the net were non-fish-eye transformed images. For all four datasets we used (a cell culture, two tissue sections and a dataset containing images of animals), training with fisheye-transformed images resulted in significantly higher accuracies. Though our method is robust, it is easy and fast to use. Furthermore, the new fisheye distortion approach is generally applicable to any kind of image data, where the environment has an influential role, as demonstrated by applying this method on the iWildCam2020 dataset.

The highest improvement in accuracy appeared when we applied our method on tissue section images. Generally, microenvironmental differences are visible in tissue histology studies. These are obvious manifestations of the cooperation and interdependence of different cells, which are also characterised histoanatomically. For example, the robust adjacency information of endothelial cells is the presence of a lumen on one side of them, or the presence of well-known restricted cell types such as connective tissue cells, smooth muscle cells on the other side.

In the case of cell cultures, we also achieved an improvement in phenotyping accuracy. The explanation relies on two factors. Firstly, homogeneous-looking areas do not consist of molecularly completely identical cells^24^. This is captured by the fisheye transformation, similarly as described in case of cancer tissues. Secondly, in homogenous cellular regions the neighbourhood may provide a statistically more stable and consequently more powerful basis for decision.

We observed minor improvement in accuracy in the case of iWildCam data. A possible explanation is that during the recording of this dataset, fixed positioned cameras were used. Because of this, depending on whether the animals are positioned near or far from the camera, one can see larger, smaller or no surroundings. Therefore, increasing the window size, it may occur that we do not gain more information about the environment.

In conclusion, we show that the incorporation of the microenvironment into machine-based decisions can improve the task of classifying single cells into phenotypic classes. This confirms the fact that cellular structures are not arbitrarily organised and it is beneficial taking these macro structures into consideration. We also show that using a non-uniformly sampling of the original image data for deep learning training and inference is feasible and can further improve accuracy. A potential extension to the presented approach could rely on the introduction of data transformer layers that are capable of learning non-linear spatial sampling functions.

## Data Availability

The image data and machine learning training sets generated and analysed during the current study are available from the corresponding author on request.

## Code Availability

The code for the fisheye transformation is available at: https://bitbucket.org/biomag/fisheye-transformation/src/master/

The Matlab code for transfer learning is available at: https://github.com/timitoth/transferlearning_forfisheye

## Ethical authorization

Ethics oversight: University of Szeged Ethics Committee.

## Acknowledgements

T.T., D.B., and P.H. acknowledge support from the LENDULET-BIOMAG Grant (2018-342), from OTKA-SNN, from TKP2021-EGA09, from H2020-COMPASS-ERAPerMed, from CZI Deep Visual Proteomics, from H2020-DiscovAir, H2020-Fair-CHARM and from the ELKH-Excellence grant. The authors thank Dora Bokor, PharmD (Szeged, Hungary), for proofreading the manuscript.

## Author contributions

P.H. conceived and led the project. F.S. co-supervised the project. T.T. designed the pipeline.

D.B. implemented the fisheye transformations in python. F.K. helped in the annotation. T.T. prepared the figures. T.T., P.H. and F.S. designed the experiments and analysed the data. All authors read and approved the final manuscript.

## References

1. Toth, T., Balassa, T., Bara, N., Kovacs, F. & Kriston, A. Environmental properties of cells improve machine learning-based phenotype recognition accuracy. Sci. Rep. 1–9 (2018) doi:10.1038/s41598-018-28482-y.

2. Wang, X. et al. Three-dimensional intact-tissue sequencing of single-cell transcriptional states. Science 361, (2018).

3. Zhu, Q., Shah, S., Dries, R., Cai, L. & Yuan, G.-C. Identification of spatially associated subpopulations by combining scRNAseq and sequential fluorescence in situ hybridization data. Nat. Biotechnol. 36, 1183–1190 (2018).

4. Standke, S. J., Colby, D. H., Bensen, R. C., Burgett, A. W. G. & Yang, Z. Mass Spectrometry Measurement of Single Suspended Cells Using a Combined Cell Manipulation System and a Single-Probe Device. Anal. Chem. 91, 1738–1742 (2019).

5. Lock, J. G. & Stromblad, S. Systems microscopy: an emerging strategy for the life sciences. Exp. Cell Res. 316, 1438–1444 (2010).

6. Meijering, E., Carpenter, A. E., Peng, H., Hamprecht, F. A. & Olivo-Marin, J.-C. Imagining the future of bioimage analysis. Nat. Biotechnol. 34, 1250–1255 (2016).

7. Caicedo, J. C. et al. Data-analysis strategies for image-based cell profiling. Nat. Methods 14, 849–863 (2017).

8. Dufour, A. C., Jonker, A. H. & Olivo-Marin, J. C. Deciphering tissue morphodynamics using bioimage informatics. Philos. Trans. R. Soc. B Biol. Sci. 372, (2017).

9. Keller, P. J. Imaging Morphogenesis: Technological Advances and Biological Insights. Science (80-.). 340, 1234168 (2013).

10. Grys, B. T. et al. Machine learning and computer vision approaches for phenotypic profiling. J. Cell Biol. 216, 65–71 (2016).

11. Scheeder, C., Heigwer, F. & Boutros, M. Machine learning and image-based profiling in drug discovery. Curr. Opin. Syst. Biol. 10, 43–52 (2018).

12. Lin, D., Sun, L., Toh, K.-A., Zhang, J. B. & Lin, Z. Biomedical image classification based on a cascade of an SVM with a reject option and subspace analysis. Comput. Biol. Med. 96, 128–140 (2018).

13. Molnar, C. et al. Accurate Morphology Preserving Segmentation of Overlapping Cells based on Active Contours. Sci. Rep. 6, 32412 (2016).

14. Meijering, E., Dzyubachyk, O., Smal, I. & van Cappellen, W. A. Tracking in cell and developmental biology. Semin. Cell Dev. Biol. 20, 894–902 (2009).

15. Pratapa, A., Doron, M. & Caicedo, J. C. Image-based cell phenotyping with deep learning. Curr. Opin. Chem. Biol. 65, 9–17 (2021).

16. Gupta, A. et al. Deep Learning in Image Cytometry: A Review. Cytom. Part A 95, 366–380 (2019).

17. Moen, E. et al. Deep learning for cellular image analysis. Nat. Methods 16, 1233–1246 (2019).

18. Mattiazzi Usaj, M. et al. Systematic genetics and single-cell imaging reveal widespread morphological pleiotropy and cell-to-cell variability. Mol. Syst. Biol. 16, e9243 (2020).

19. Coudray, N. et al. Classification and mutation prediction from non–small cell lung cancer histopathology images using deep learning. Nat. Med. 24, 1559–1567 (2018).

20. Sullivan, D. P. et al. Deep learning is combined with massive-scale citizen science to improve large-scale image classification. Nat. Biotechnol. 36, 820–828 (2018).

21. Ouyang, W. et al. Analysis of the Human Protein Atlas Image Classification competition. Nat. Methods 16, 1254–1261 (2019).

22. Thul, P. J. et al. A subcellular map of the human proteome. Science (80-.). 356, eaal3321 (2017).

23. Raj, A. & van Oudenaarden, A. Nature, nurture, or chance: stochastic gene expression and its consequences. Cell 135, 216–226 (2008).

24. Snijder, B. et al. Population context determines cell-to-cell variability in endocytosis and virus infection. Nature 461, 520–523 (2009).

25. Bove, A. et al. Local cellular neighborhood controls proliferation in cell competition. Mol. Biol. Cell 28, 3215–3228 (2017).

26. Mesa, K. R. et al. Homeostatic Epidermal Stem Cell Self-Renewal Is Driven by Local Differentiation. Cell Stem Cell 23, 677-686.e4 (2018).

27. Shaya, O. et al. Cell-Cell Contact Area Affects Notch Signaling and Notch-Dependent Patterning. Dev. Cell 40, 505-511.e6 (2017).

28. Sahin, C. The Geometry and Usage of the Supplementary Fisheye Lenses in Smartphones. Smartphones from an Appl. Res. Perspect. (2017) doi:10.5772/intechopen.69691.

29. Schmalstieg, D. & Hollerer, T. Augmented reality: principles and practice. (Addison-Wesley Professional, 2016).

30. Sáez, Á. et al. Real-Time Semantic Segmentation for Fisheye Urban Driving Images Based on ERFNet. Sensors vol. 19 (2019).

31. Tseng, D., Chen, C. & Tseng, C. Automatic detection and tracking in multi-fisheye cameras surveillance system. Int. J. Comput. Elect. Eng. 9, (2017).

32. Li, T., Tong, G., Tang, H., Li, B. & Chen, B. FisheyeDet: A Self-Study and Contour-Based Object Detector in Fisheye Images. IEEE Access 8, 71739–71751 (2020).

33. Silberstein, S., Levi, D., Kogan, V. & Gazit, R. Vision-based pedestrian detection for rear-view cameras. In 2014 IEEE Intelligent Vehicles Symposium Proceedings 853–860 (2014). doi:10.1109/IVS.2014.6856399.

34. Bertozzi, M., Castangia, L., Cattani, S., Prioletti, A. & Versari, P. 360° Detection and tracking algorithm of both pedestrian and vehicle using fisheye images. In 2015 IEEE Intelligent Vehicles Symposium (IV) 132–137 (2015). doi:10.1109/IVS.2015.7225675.

35. Jaderberg, M., Simonyan, K., Zisserman, A. & kavukcuoglu, koray. Spatial Transformer Networks. In Advances in Neural Information Processing Systems (eds. Cortes, C., Lawrence, N., Lee, D., Sugiyama, M. & Garnett, R.) vol. 28 (Curran Associates, Inc., 2015).

36. Caie, P. D. et al. High-Content Phenotypic Profiling of Drug Response Signatures across Distinct Cancer Cells. Mol. Cancer Ther. 9, 1913 LP – 1926 (2010).

37. Piccinini, F. et al. Advanced Cell Classifier: User-Friendly Machine-Learning-Based Software for Discovering Phenotypes in High-Content Imaging Data. Cell Syst. 4, 651-655.e5 (2017).

38. Mund, A. et al. AI-driven Deep Visual Proteomics defines cell identity and heterogeneity. bioRxiv (2021) doi:10.1101/2021.01.25.427969.

39. Achanta, R. et al. SLIC Superpixels Compared to State-of-the-Art Superpixel Methods. IEEE Trans. Pattern Anal. Mach. Intell. 34, 2274–2282 (2012).

