## Supplementary for "Show me your neighbour and I tell what you are: fisheye transformation for deep learning-based single-cell phenotyping"

### *Section 1 - Fisheye transformation*

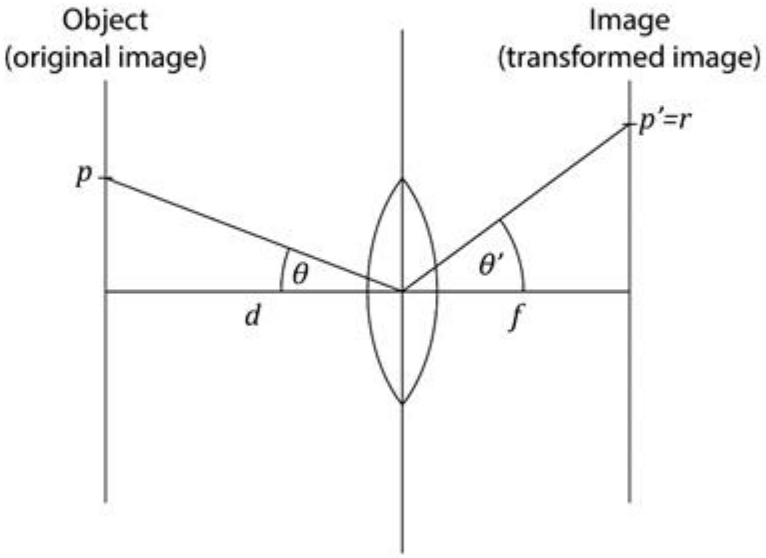


In fisheye transformation, the position of the projection of a given real world point can be determined from the angle of the incident ray. This can be calculated using a mapping function:

$$r = m(f, \theta),$$

where $f$ is the focal length. For a given $f$ we can reformulate the above equation as

$$r = m_f(\theta).$$

The mapping function is an inherent function to fisheye lenses. Possible mapping functions are listed in Supplementary Section 2. All of these functions are invertible:

$$\theta = m_f^{-1}(r).$$

The following relation is also valid for $\theta$:

$$tan\,\theta\ = \frac{p}{d},$$

where $d$ is the distance between point $p$ and the centre of the lens, measured along the axis of the lens. In our case the object points are the pixels of an image, and $d$ is constant across the whole image.

In image transformation tasks the transformation function is usually given as an inverse mapping, which provides the source position of each output pixel, in this case, the value of $p$ for each output position $r$. From the equations above, *p* can be calculated as follows:

$$p = d \cdot tan\, m_f^{-1}(r)\,.$$

This equation contains two free parameters: $d$ and $f$. The value of $d$ appears as a scalar multiplier, hence it affects magnification only. The value of $f$ affects both the scale and the magnitude of distortion. However, in our case we endeavoured to change the strength of the fisheye effect of the transformation only, leaving the scale of the selected area unaffected. In order to achieve the desired effect, it is possible to connect the value of $d$ to the value of $f$ in such a way that it unaffects scaling.

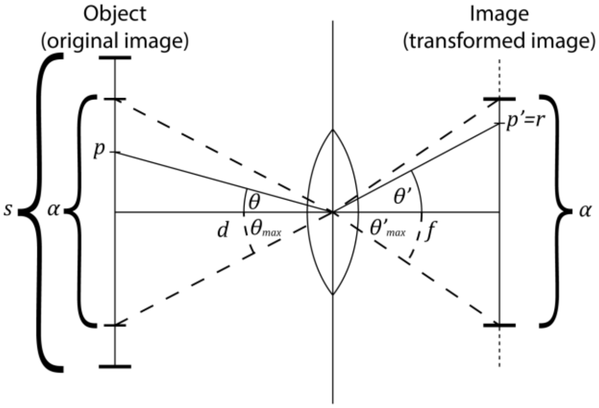


Let's mark the size of the selected area with $\alpha$. We wish to keep the position of the corner points intact while applying the fisheye distortion. In this case a new equation is introduced as follows:

$$tan\, \theta_{max} = \frac{\alpha}{2d},$$

where $\theta_{max}$ is the angle of the incoming ray from the borders of the selected area. As for the border points are expected to be transformed into themselves,

$$\theta_{max} = m_f^{-1}\left(\frac{\alpha}{2}\right)$$

is also valid. Combining these two equations gives

$$d = \frac{\alpha}{2\, tan\ m_f^{-1}\left(\frac{\alpha}{2}\right)}.$$

It is evident that the last equation is determined by $f$ as a modifiable parameter (when the value of $\alpha$ is fixed). Thus, the final form of the equation for the fisheye transformation is

$$p = \frac{\alpha}{2} \cdot \frac{tan\ m_f^{-1}(r)}{tan\ m_f^{-1}\left(\frac{s}{2}\right)}.$$

As an easy-to-read interpretation, we normalise the value of θ to the range of $[0, 1]$, and then rescale it to the range $\left[0, \frac{\alpha}{2}\right]$.

Note that in the calculations above it is assumed that the selected area of interest is in the middle of the image. However, with a simple translation, the calculations are valid for any arbitrary image positions.

### *Section 2 - Mapping functions*

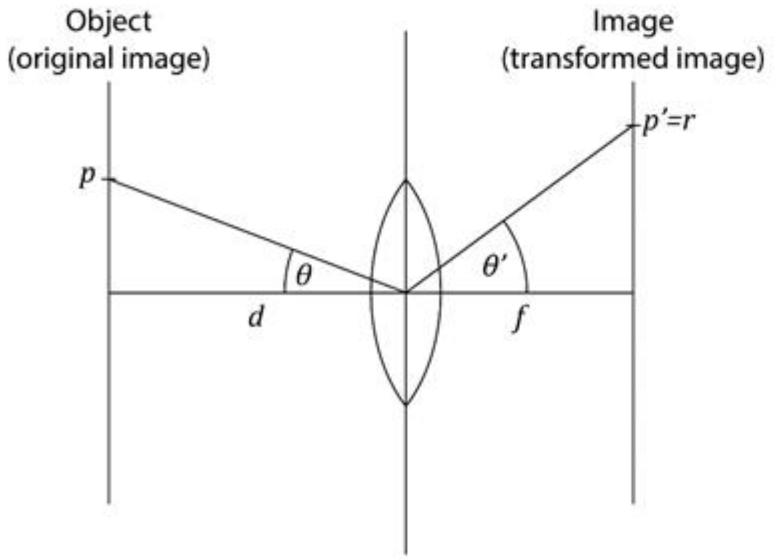


In cameras with wide-angle lenses, the object is located in the image according to the mapping function of the lens. The mapping function defines the position of the object from the centre of the image (*r*) as a function of the focal distance (*f*) and the angle from the optical axis (θ).

The functions in wide-angle lens cameras include the following:

- Rectilinear:

$$r = f \tan \theta$$

- Fisheye
    - Equidistant

$$r = f\theta$$

    - Equisolid angle

$$r = 2f \sin \frac{\theta}{2}$$

    - Stereographic

$$r = 2f\tan\frac{\theta}{2}$$

- Orthographic

$$r = f\sin\theta$$

***Section 3 - Deep learning parameters***

MiniBatchSize: 64

MaxEpochs: 100

InitialLearnRate: 3e-4

LearnRateDropFactor: 0.3

LearnRateDropPeriod: 50

Shuffle: every epoch

The whole code for the deep learning can be found in Supplementary Data1.

**Section 4 – *Significance tests***

*MCF-7*

Our best result (accuracy: 91.38%) for this dataset with the fisheye-transformation appears with a window size of 543 pixels with a focal length of 130 arbitrary units when we used ResNet50. We compare this result to a baseline, where we used ResNet50 (accuracy: 84.31%). For the baseline we cropped out images around the cells' centre with 192x192 pixel diameter (so in this case, we haven't performed fisheye transformation on the original images). Deep learning calculations were run 5-5 times on both the baseline and the fisheye transformed data.

| MCF7 baseline accuracies | MCF7 best accuracies |
|---|---|
| 82,23 | 92 |
| 84,89 | 90,15 |
| 85,03 | 91,08 |
| 84,17 | 91,68 |
| 85,23 | 91,99 |

As a statistical procedure, two-sample t-test was performed, we considered the result significant at $p < 0.05$.

| ***Descriptive statistics*** | | N | Mean | SD | SEM | Median |
|---|---|---|---|---|---|---|
| MCF7_baseline | | 5 | 84,31 | 1,22955 | 0,54987 | 84,89 |
| MCF7_best | | 5 | 91,38 | 0,78253 | 0,34996 | 91,68 |
| | *Difference* | 5 | -7,07 | | 0,65179 | -6,76 |

| ***t-Test Statistics*** | t Statistic | DF | Prob>\|t\| |
|---|---|---|---|
| Equal Variance Assumed | -10,8471 | 8 | 4,61E-06 |
| Equal Variance NOT Assumed (Welch Correction) | -10,8471 | 6,78368 | 1,56E-05 |

Null hypothesis: mean1-mean2 = 0

Alternative hypothesis: mean1-mean2 <> 0

<u>At the 0.05 level, the 7.07% difference in accuracy is significant.</u>

*Urinary Bladder Cancer*

Our best result (accuracy: 98.14%)for this dataset with the fisheye-transformation appears with a window size of 272 pixels with a focal length of 150 arbitrary units when we used ResNet50. We compare this result to a baseline, where we used ResNet50 (accuracy: 94.41%). For the baseline we cropped out images around the cells' centre with 192x192 pixel diameter (so in this case, we haven't performed fisheye transformation on the original images). Deep learning calculations were run 5-5 times on both the baseline and the fisheye transformed data.

| UBC baseline accuracies | UBC best accuracies |
|---|---|
| 94,12 | 97,94 |
| 94,68 | 98,24 |
| 94,04 | 98,24 |
| 94,85 | 97,97 |
| 94,36 | 98,31 |

As a statistical procedure, two-sample t-test was performed, we considered the result significant at $p < 0.05$.

| ***Descriptive statistics*** | | N | Mean | SD | SEM | Median |
|---|---|---|---|---|---|---|
| UBC_baseline | | 5 | 94,41 | 0,35 | 0,15652 | 94,36 |
| UBC_best | | 5 | 98,14 | 0,17161 | 0,07675 | 98,24 |
| | *Difference* | 5 | -3,73 | | 0,17433 | -3,82 |

| ***t-Test Statistics*** | t Statistic | DF | Prob>\|t\| |
|---|---|---|---|
| Equal Variance Assumed | -21,3965 | 8 | 2,40E-08 |
| Equal Variance NOT Assumed (Welch Correction) | -21,3965 | 5,81818 | 9,37E-07 |

Null hypothesis: mean1-mean2 = 0

Alternative hypothesis: mean1-mean2 <> 0

<u>At the 0.05 level, the 3.83% difference in accuracy is significant.</u>

*Lung Cancer*

Our best result (accuracy: 99.46%)for this dataset with the fisheye-transformation appears with a window size of 272 pixels with a focal length of 170 arbitrary units when we used inceptionV3. We compare this result to a baseline, where we used inceptionV3 (accuracy: 97.25%). For the baseline we cropped out images around the cells' centre with 192x192 pixel diameter (so in this case, we haven't performed fisheye transformation on the original images). Deep learning calculations were run 5-5 times on both the baseline and the fisheye transformed data.

| LC baseline accuracies | LC best accuracies |
|---|---|
| 97,32 | 99,64 |
| 97,54 | 98,97 |
| 97,01 | 99,57 |
| 97,03 | 99,54 |
| 97,35 | 99,58 |

As a statistical procedure, two-sample t-test was performed, we considered the result significant at $p < 0.05$.

| ***Descriptive statistics*** | | N | Mean | SD | SEM | Median |
|---|---|---|---|---|---|---|
| Lung_baseline | | 5 | 97,25 | 0,22638 | 0,10124 | 97,32 |
| Lung_best | | 5 | 99,46 | 0,27632 | 0,12357 | 99,57 |
| | *Difference* | 5 | -2,21 | | 0,15975 | -2,32 |

| ***t-Test Statistics*** | t Statistic | DF | Prob>\|t\| |
|---|---|---|---|
| Equal Variance Assumed | -13,8341 | 8 | 7,20E-07 |
| Equal Variance NOT Assumed (Welch Correction) | -13,8341 | 7,70198 | 1,03E-06 |

Null hypothesis: mean1-mean2 = 0

Alternative hypothesis: mean1-mean2 <> 0

At the 0.05 level, the 2.21% difference in accuracy is significant.

*iWildCam2020*

Our best result (accuracy: 95.48%)for this dataset with the fisheye-transformation appears when 2.5× the size of bounding boxes were considered as the neighbourhood feature and focal length was set to 150 units and we used ResNet50. We compare this result to a baseline, where we used ResNet50 (accuracy: 95.3%). For the baseline we cropped out images with different dimensions based on the bounding boxes provided by the Kaggle organisers (so in this case, we haven't performed fisheye transformation on the images). Deep learning calculations were run 5-5 times on both the baseline and the fisheye transformed data.

| iWildCam baseline accuracies | iWildCam best accuracies |
|---|---|
| 95,32 | 95,51 |
| 95,29 | 95,46 |
| 95,27 | 95,49 |
| 95,33 | 95,46 |
| 95,29 | 95,48 |

As a statistical procedure, two-sample t-test was performed, we considered the result significant at p <0.05.

| ***Descriptive statistics*** | | N | Mean | SD | SEM | Median |
|---|---|---|---|---|---|---|
| iWildCam_baseline | | 5 | 95,3 | 0,02449 | 0,01095 | 95,29 |
| iWildCam_best | | 5 | 95,48 | 0,02121 | 0,00949 | 95,48 |
| | *Difference* | 5 | -0,18 | | 0,01449 | -0,19 |

| ***t-Test Statistics*** | t Statistic | DF | Prob>\|t\| |
|---|---|---|---|
| Equal Variance Assumed | -12,4212 | 8 | 1,65E-06 |
| Equal Variance NOT Assumed (Welch Correction) | -12,4212 | 7,84 | 1,96E-06 |

Null hypothesis: mean1-mean2 = 0

Alternative hypothesis: mean1-mean2 <> 0

At the 0.05 level, the 0.18% difference in accuracy is significant.

**Section 5 – *Classification in the MCF-7 dataset based on nuclei and cytoplasm***

Looking at the fisheye transformed images, it is reasonable to wonder whether we can achieve similarly high results by using only the nuclei to train our deep-learning network. In the case of the MCF-7 dataset we did nuclei and cytoplasm segmentation (as described in Materials and Methods). To demonstrate that the increase in classification accuracy is due to the inclusion of the environment and the fisheye transformation (as we concluded in our paper), we ran calculations on images showing only the nucleus, or the nucleus and cytoplasm of the cell. We used the same ResNet50 deep-learning network as in the manuscript for the baseline and fisheye calculations (we have got the best results with this network). The results show the same tendency that we expected, that the classification accuracy is lowest when the network sees only the nuclei and best when we use the fisheye transformed images.

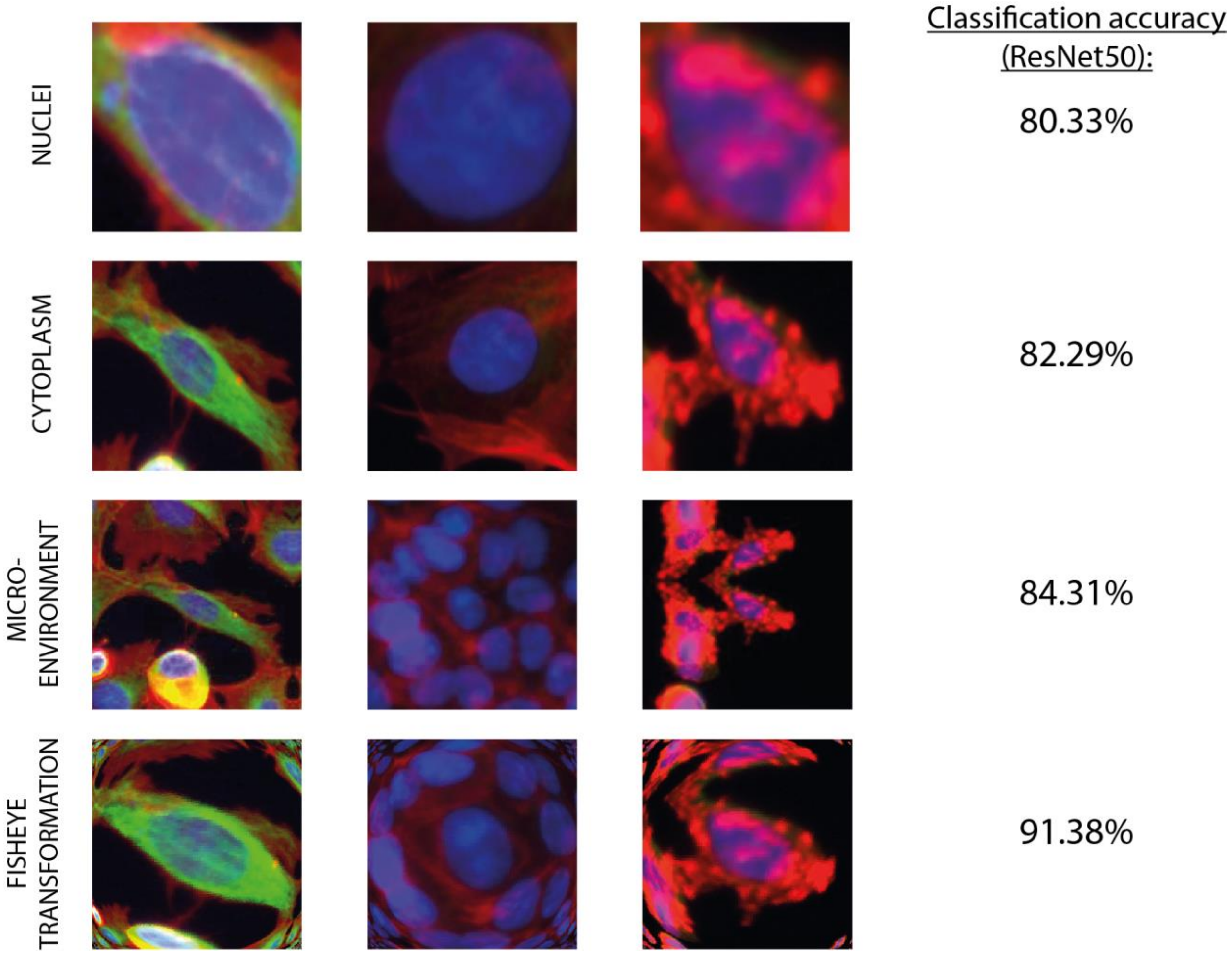

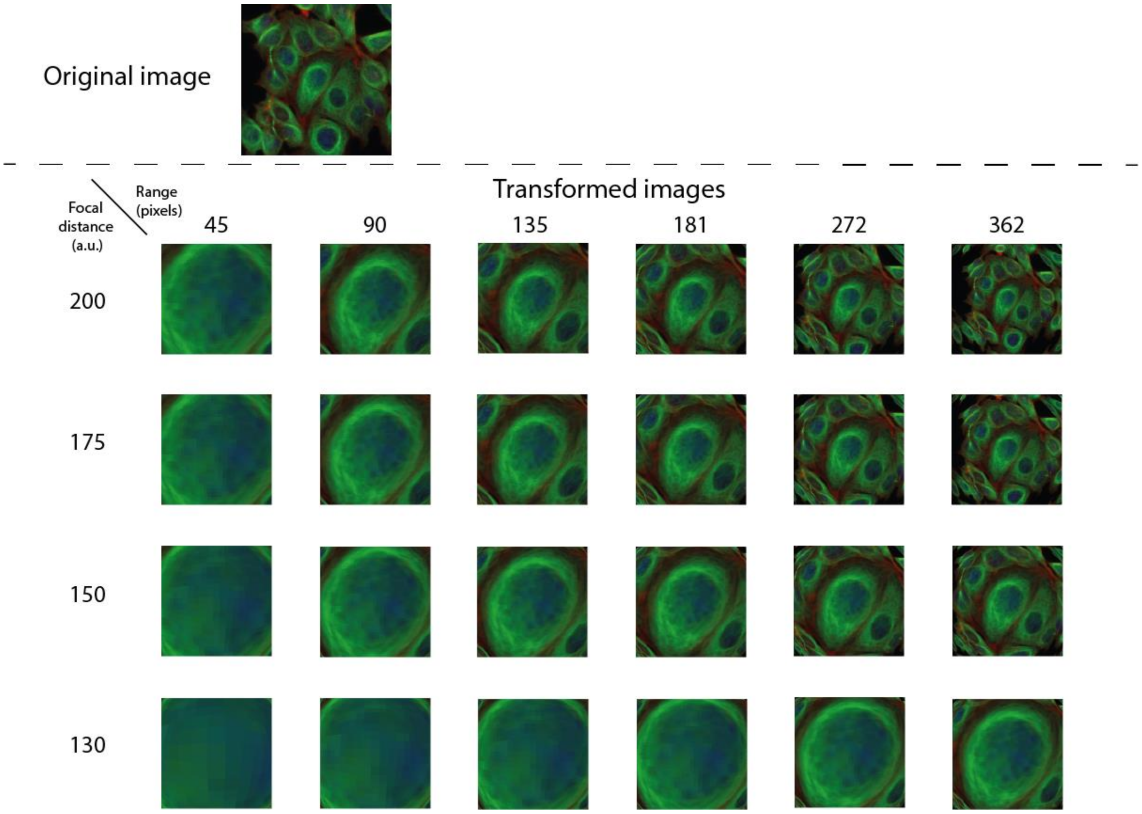


**Figure S1.** Related to Figure 1. Examples of the fisheye transformation with different window sizes (ranges) and focal distances using the 'equidistant' mapping function in the MCF-7 cell culture dataset.

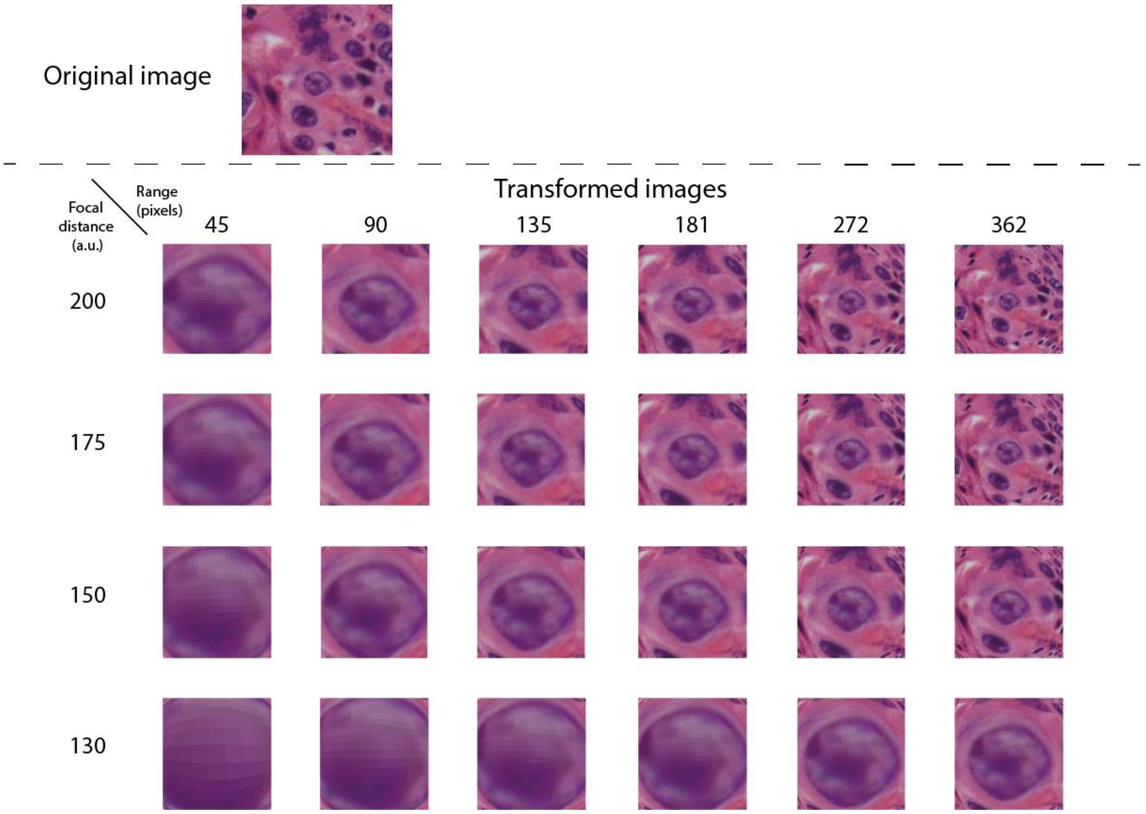


**Figure S2.** Related to Figure 1. Examples of the fisheye transformation with different window sizes (ranges) and focal distances using the 'equidistant' mapping function in the urinary bladder cancer tissue dataset.

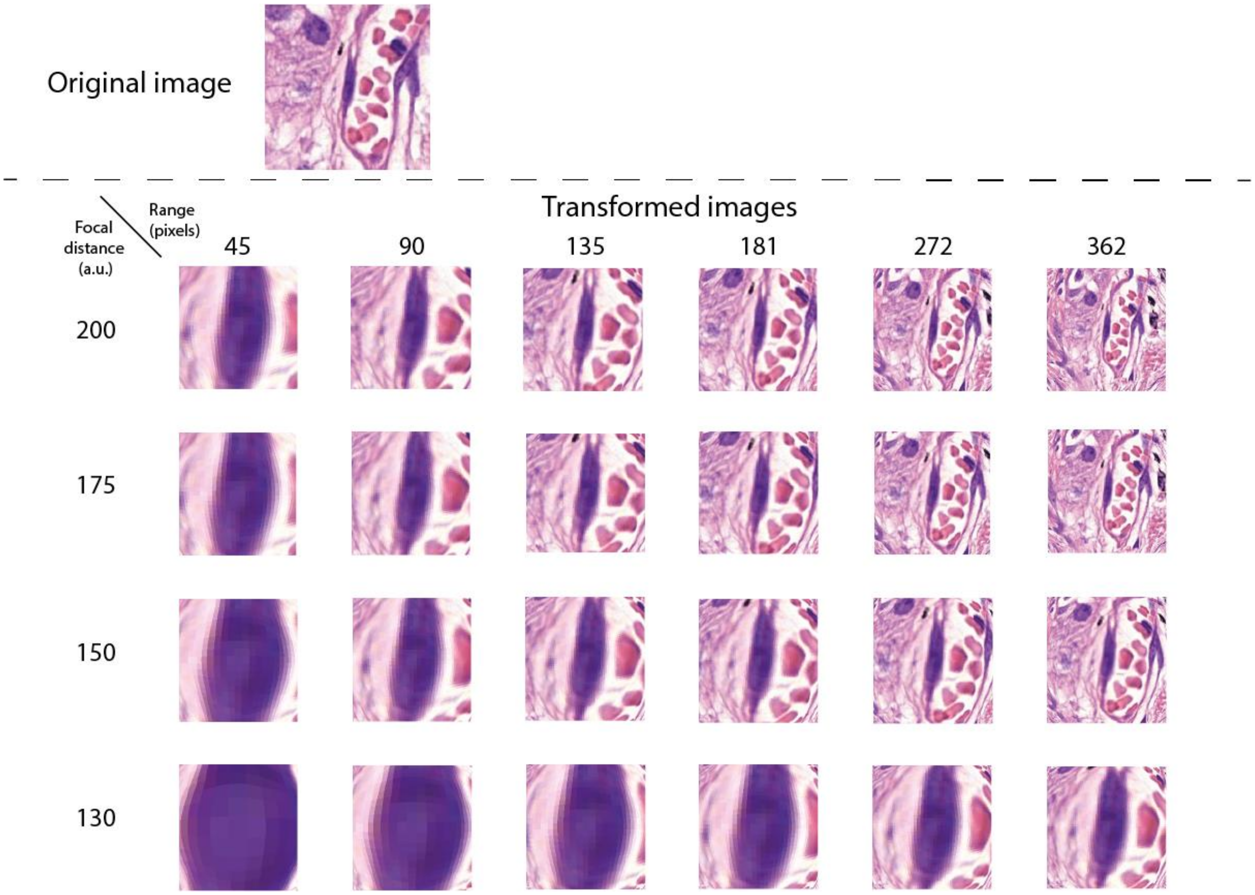


**Figure S3.** Related to Figure 1. Examples of the fisheye transformation with different window sizes (ranges) and focal distances using the ‘equidistant’ mapping function in the lung cancer tissue dataset.

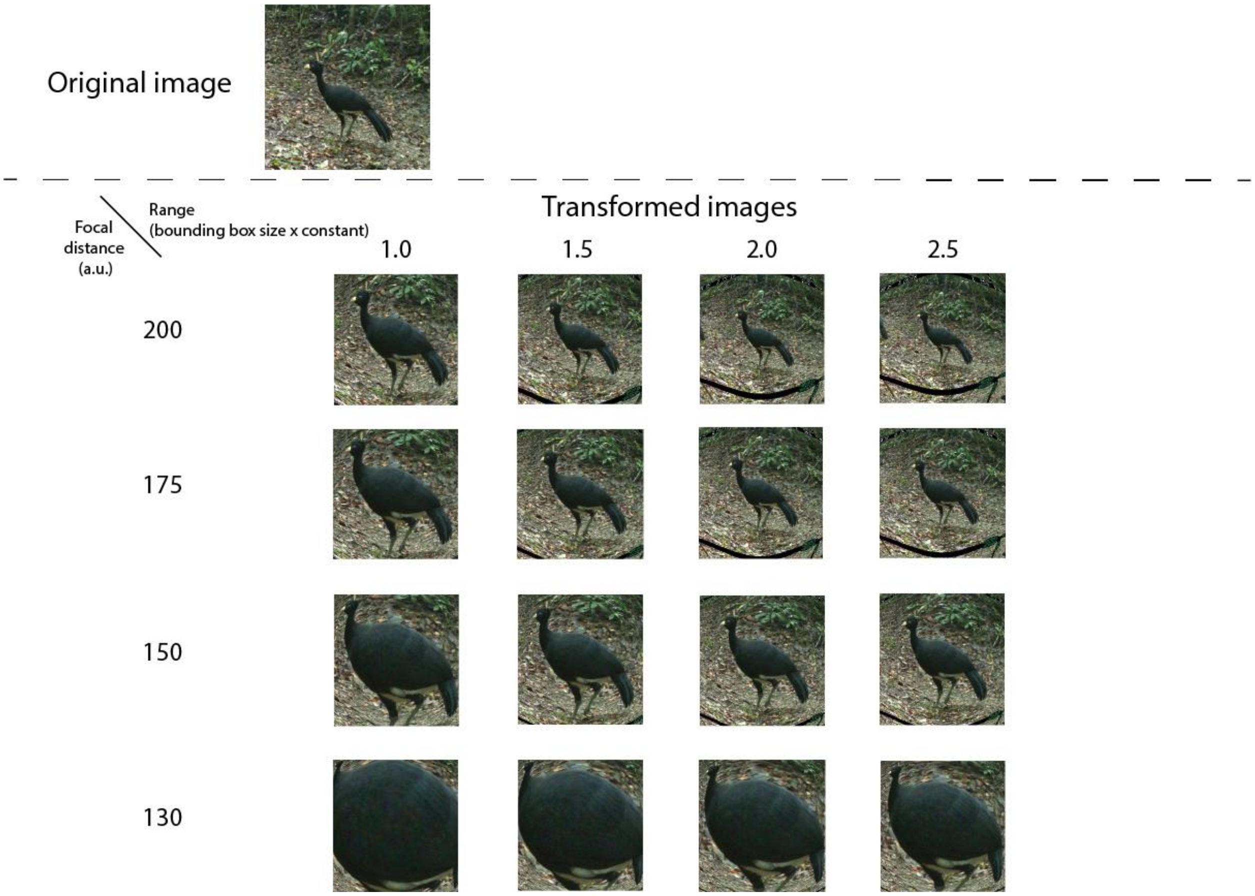


**Figure S4.** Related to Figure 1. Examples of the fisheye transformation with different window sizes (ranges) and focal distances using the 'equidistant' mapping function in the iWildCam2020 dataset.
